# AASRA: An Anchor Alignment-Based Small RNA Annotation Pipeline

**DOI:** 10.1101/132928

**Authors:** Chong Tang, Yeming Xie, Wei Yan

## Abstract

SncRNA-Seq has become a routine for sncRNA profiling; however, software packages currently available are either exclusively for miRNA or piRNA annotation (e.g., miRDeep, miRanalyzer, Shortstack, PIANO), or for direct mapping of the sequence reads to the genome (e.g., Bowtie 2, SOAP and BWA), which tend to generate inaccurate counting due to repetitive matches to the genome or sncRNA homologs. Moreover, novel sncRNA variants in the sequencing reads, including those bearing small overhangs or internal insertions, deletions or mutations, are totally excluded from counting by these algorithms, leading to potential quantification bias. To overcome these problems, a comprehensive software package that can annotate all known small RNA species with adjustable tolerance towards small mismatches is needed. AASRA is based on our unique anchor alignment algorithm, which not only avoids repetitive or ambiguous counting, but also distinguishes mature miRNA from precursor miRNA reads. Compared to all existing pipelines for small RNA annotation, AASRA is superior in the following aspects: 1) AASRA can annotate all known sncRNA species simultaneously with the capability of distinguishing mature and precursor miRNAs; 2) AASRA can identify and allow for inclusion of sncRNA variants with small overhangs and/or internal insertions/deletions into the final counts; 3) AASRA is the fastest among all small RNA annotation pipelines tested. AASRA represents an all-in-one sncRNA annotation pipeline, which allows for high-speed, simultaneous annotation of all known sncRNA species with the capability to distinguish mature from precursor miRNAs, and to identify novel sncRNA variants in the sncRNA-Seq sequencing reads.

**Availability and Implementation:** The AASRA software is freely available at <u>https://github.com/biogramming/AASRA.</u>

## 1. Introduction

Given their critical regulatory roles, small noncoding RNAs (sncRNAs) have become a major focus in biomedical research [1, 2]. The next-gen sequencing technologies have allowed for the identification of hundreds thousands of sncRNAs, which have been categorized into many unique sncRNA species, e.g., microRNAs (miRNAs) [3-5], endogenous small interference RNAs (endo-siRNAs) [6, 7], PIWI-interacting RNAs (piRNAs) [8-11], small nucleolar RNAs (snoRNAs) [12], tRNA-derived small RNAs (tsRNAs) [13, 14], mitochondrial genome –encoded small RNAs (mitosRNAs) [15], etc. Among these sncRNAs, miRNAs and piRNAs have been studied extensively for the past decade largely because they were discovered first [8-11, 16]. To help investigators identify known and to predict novel miRNAs or piRNAs based on sncRNA next-gen sequencing (sncRNA-Seq) data, several software packages have been developed, e.g., ShortStack [17], miRanalyzer [18], miRDeep [19], PIANO [20], etc. Using these pipelines, researchers have not only validated previously reported sncRNAs, but also predicted sncRNAs based on their unique structural (e.g., length, stem-loop structure, etc.) and genomic features (e.g., repetitive sequences). Currently, there are many sncRNA databases, e.g., miRBase [21], piRNABank [22], piRNA Cluster Database [23], Rfam [24-26], snoRNA-LBME-db [27], etc., where known and predicted sncRNAs (for some of the databases) are collected. These databases serve as important resources because investigators can download these sncRNAs and use them as reference sequences to annotate their own sncRNA-Seq data for sncRNA identification and quantitation. Currently, one way to annotate sncRNAs is to map the sncRNA-Seq reads directly to the reference genome using Bowtie [28], SOAP [29] or BWA [30], followed by counting based on the genome feature file (e.g., GFF/GTF). Alternatively, the sncRNA-Seq reads can be aligned to the reference sncRNA sequences downloaded from the available databases, using sequence alignment software packages, e.g., miRDeep [19, 31]. Methods based on alignment to both the reference sncRNAs and the genome have also been developed, e.g., miRanalyzer [18, 32]. While these pipelines perform well when used for annotating sncRNAs that are already collected in the databases, they can neither distinguish between mature and precursor miRNAs, nor count sncRNA variants with small overhangs and/or internal insertions, deletions or mutations. In addition, there are no software packages that allow for simultaneous annotation of all known small RNA species. To overcome these problems, we developed a new software package, which we named “AASRA” (for <u>A</u>nchor <u>A</u>lignment-based <u>S</u>mall <u>R</u>NA <u>A</u>nnotation). AASRA is based on a novel alignment algorithm and can annotate sncRNAs of all known species collected in various sncRNA databases with a much higher mapping rate and accuracy, as well as speed, compared to all existing software packages currently available for sncRNA annotation.

## 2. Materials and Methods

### 2.1 Small noncoding RNA reference data

The reference sncRNA datasets consists of mature and precursor miRNAs in the miRBase (release 21) [21], tRNAs in the Genomic tRNA Database [33], piRNAs in the piRNABank [22] and piRNA Cluster Database [23], rRNAs, snoRNAs, snRNAs and mitochondrial RNAs in ENSEMBL (release 76) [34-36], and endo-siRNAs in DeepBase [37].

### 2.2 Simulation data

Simulation sequences were based upon sncRNA sequences from the known sncRNA databases. sncRNA variant sequences, including 1-2nt overhangs, internal insertions, deletions and mutations, were generated by randomly adding or changing 1-2nts at either end or internally using R script of the Biostrings package. To generate the simulation Fasta file, individual sncRNAs were randomly duplicated such that the counts for each ranged from 1 to 50.

### 2.3 Anchor alignment

Anchor sequences (5-10bp) were added to both ends of the reference sncRNAs and the sequencing reads, as well as simulation sequences using the Python script. “Bowtie2-build” was employed to index all the anchored reference sncRNAs. The anchored sequencing reads/simulation sequences were then aligned to the indexed anchored reference sncRNAs using Bowtie2 [38]. The FeatureCounts [39] was used to summarize the counts in the alignment file. The same procedure was used to align the non-anchored sequencing reads or simulation sequences to the indexed, non-anchored reference sncRNA sequences.

### 2.4 Genome alignment

Bowtie2-build was used to index the mouse genome (NCBI_Assembly: GCA_000001635.2). The sequencing data were aligned to the indexed genome using Bowtie 2. The FeatureCount was used to summarize the reads in the alignment file based on mmu.gff3 (miRbase V21).

### 2.5 miRNA annotation using miRDeep

The GRCm38 mouse genome was built according to the user manual of miRDeep [19, 31]. Both miRNA sequencing reads/simulation dataset and GRCm38 pre-built genome were loaded for alignment analyses using the default setting of miRDeep. Scatter plots were generated to correlate the predicted counts (by miRDeep) with the standard counts (simulation counts).

### 2.6 miRNA annotation using ShortStack

“Bowtie2-build” was used to generate the indexed mouse genome (NCBI_Assembly: GCA_000001635.2). The simulation data were then aligned to the indexed genome using Bowtie 2 (ShortStack --readfile --outdir --genomefile), and the hits were --locifile --outdir --genomefile). Scatter plots were generated to correlate the predicted counts (by ShortStack) with the standard counts (simulation counts).

### 2.7 miRNA annotation using miRanalyzer

The stand-alone version of miRanalyzer was downloaded and installed according to the user manual [18, 32]. The pre-built, Bowtie2-indexed genome sequences (UCSC mm9) were used as the reference mouse genome in miRanalyzer. The mature and precursor miRNA sequences were used as the sncRNA reference dataset. miRNA simulation data with or without overhangs were analyzed using the default parameters. Scatter plots were generated to correlate the predicted counts (by miRanalyzer) with the standard counts (simulation counts).

### 2.8 Mouse sperm sncRNA-Seq

The Institutional Animal Care and Use Committee (IACUC) of the University of Nevada, Reno approved the use of mice (Protocol#00494) for sperm collection and sncRNA-Seq. Mouse epididymal sperms were collected in the HEPES-HTF medium, and a “swim-up” procedure was performed so that only motile sperm were selected for sncRNA-Seq [40]. Total RNA was isolated using the mirVana miRNA Isolation Kit (Life Technologies) following the manufacturer’s instructions. SncRNA libraries were prepared using the Ion Total RNA-Seq Kit v2 (Life Technologies), followed by sequencing using the Ion P1 chips on an Ion Proton Sequencer (Life Technologies) [40]. The sncRNA-Seq datasets have been deposited into the NCBI GEO database with the accession number of GSE81216.

### 2.9 Data management and graphics

All the data were processed using the R script and graphs were plotted using the R script of the ggplot2 package.

## 3. Results

### 3.1 The anchor alignment algorithm

The most popular sequence alignment software packages, e.g., Bowtie [28], SOAP [29] or BWA [30], are designed for mapping large RNA sequencing reads directly to the genome. However, these methods are not ideal for small RNA alignment analyses for two reasons. First, the library construction methods for large and small RNAs are fundamentally different (Figure 1A). The Illumina sequencers perform the so-called short-read sequencing, which requires shorter DNA fragments (∼200-800bp). Therefore, large RNAs have to be fragmented either physically (*via* heating or shearing) or enzymatically, followed by adaptor ligation (Figure 1A). After sequencing, the shorter reads (∼50-150nt) need to be aligned to the genome using Bowtie2-based TopHat followed by assembly using Cufflinks [41]. Fragmentation can generate numerous homologous fragments, which differ from each other by only a few nucleotides at either or both ends. Since they are all derived from the same transcripts, the downstream annotation will categorize these homologous fragments as single transcripts. In contrast, adaptors are ligated directly to small RNAs without fragmentation during sncRNA library preparation (Figure 1A) and thus, homologous fragments represent unique sncRNAs and should, therefore, be counted as individual sncRNAs. Second, mathematically, the possibility for shorter reads (∼20-40bp) to have multiple alignments in the genome is much greater, compared to that of longer reads (50-150nt); multiple mapping leads to repetitive counting during alignment, causing quantification bias (Figure 1B). A straightforward solution would be to align the sequencing reads to the corresponding sncRNA reference sequences instead of the genome. However, this direct, RNA-to-RNA mapping strategy leads to multiple alignments due to the existence of homologous sncRNAs in both the reference databases and the sequencing reads. For example, the sequencing reads of a mature miRNA would align to both the mature miRNA and its homologous precursor miRNA in the reference dataset, leading to double counting (Figure 1C). Many sncRNAs, e.g., MIWI2-bound piRNAs (i.e., pre-pachytene piRNAs), endo-siRNAs and mitosRNAs, contain a large number of homologs with only a few nucleotide differences in either or both ends (Figure 1C). Thus, one such sncRNA would align to its multiple homologs, causing repetitive counting and quantification bias (Figure 1C). Moreover, the existing alignment programs would only select the perfectly matched reads and eliminate those with minor mismatches although those may represent the sncRNAs synthesized by the cells. To overcome these problems, we developed a universal sncRNA annotation software package, AASRA, based on our unique anchor alignment algorithm (Figure 1D). AASRA first processes both the sequencing reads and the reference sequences by adding two unique anchor sequences to both ends. Then the anchored sequencing reads are aligned to the anchored sncRNA references using Bowtie 2. Finally, FeatureCounts (Subread) is used to summarize the unique read counts (Figure 1D).

**Figure 1.**
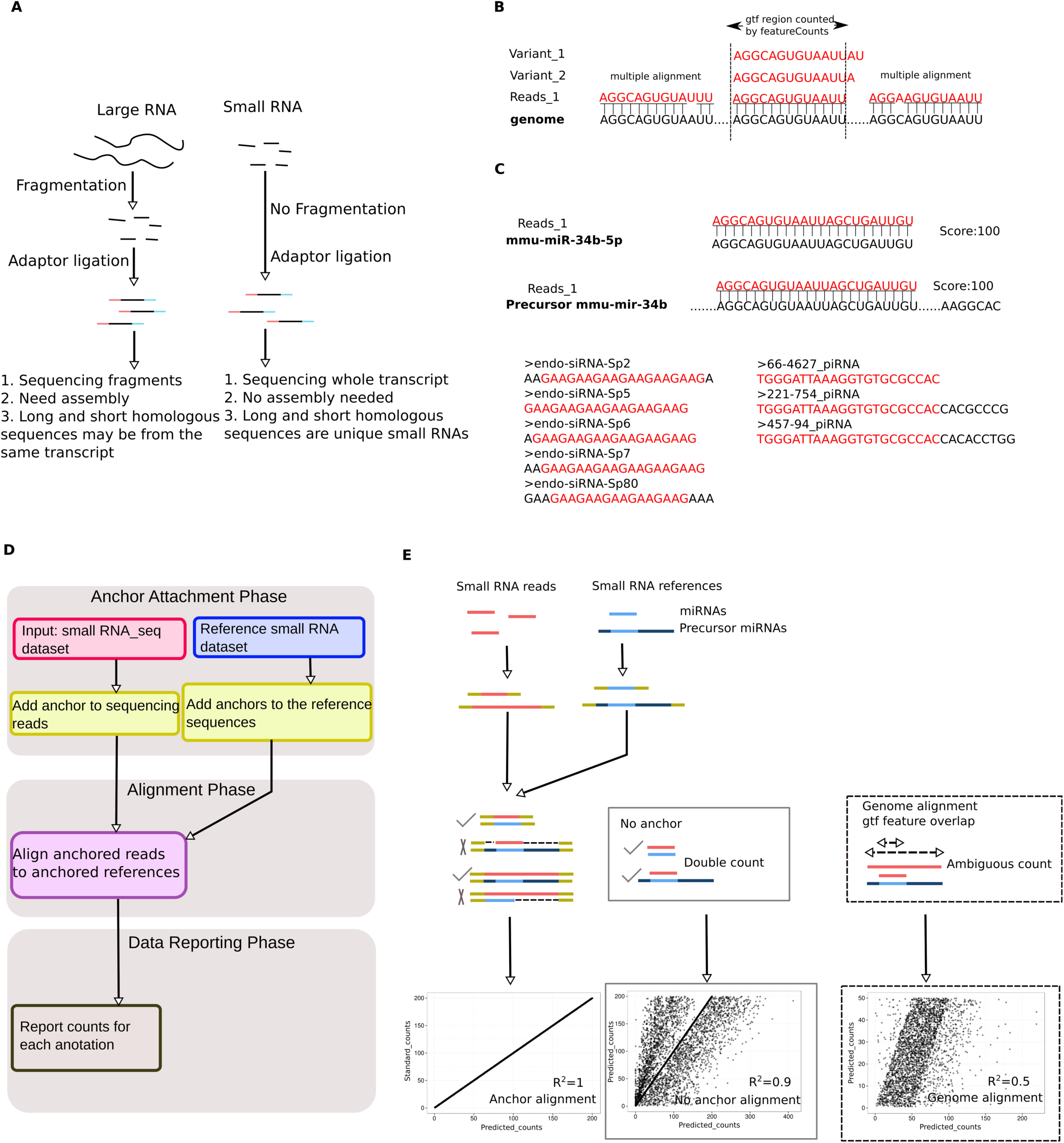
Development of the anchor alignment algorithm for sncRNA annotation. (A) Schematic illustration of the differences in large and small RNA library construction methods. Note that adaptors are directly added to the small RNAs for sncRNA-Seq, whereas fragmentation is needed before adaptor ligation for large RNA sequencing. (B) Issues associated with direct sncRNA alignment to the genome: multiple alignment of sncRNAs to the genome due to their small sizes (20-40nt), and inability to recognize sncRNA variants (e.g., homologous piRNAs, endo-siRNAs, mitosRNAs, etc.). (C) Issues associated with the direct sncRNA-sncRNA alignment algorithm: repetitive counting of mature miRNA reads (because they can be mapped to both mature and premature miRNA references), and certain sncRNA reads (e.g., endo-siRNAs and piRNAs, due to the presence of multiple, staggered sncRNA homologs in the reference databases, which differ by only several nucleotides). (D) Workflow of the anchor alignment-based sncRNA annotation (AASRA) pipeline. (E) Schematic illustration of anchor alignment algorithm. Anchors are added to both ends of the sequencing reads and the reference sncRNAs. Gap opening penalty can prevent mature miRNA sequence reads from mapping to the premature miRNA reference sequences. Perfect alignment and correct annotation of both mature and precursor miRNAs were achieved for the simulation data using the anchor alignment algorithm (*R*^*2*^=1), whereas direct alignment of the simulation data to either the sncRNA references (*R*^*2*^=0.9), or the genome (*R*^*2*^=0.5) led to partial alignment.

The anchor alignment algorithm can avoid multiple and ambiguous alignments, which are common in those straight matching algorithms (direct alignment to reference sncRNAs or to the genome by Bowtie2, or miRanalyzer, miRDeep, etc). For example, the anchored mature miRNA reads can only align to the anchored mature miRNA references. When the mature miRNA reads are aligned to the anchored reference precursor miRNAs, the gap-opening penalty would prevent double matching (Figure 1E). In this way, mature miRNAs can be readily distinguished from their corresponding precursor miRNAs during the alignment. As a proof of concept, we aligned the simulation dataset containing both mature and precursor miRNA sequences to the reference miRNA dataset downloaded from the miRBase using AASRA. The anchor alignment algorithm resulted in a perfect mapping (*R*^*2*^=1), whereas the direct alignment to the reference miRNAs or to the genome led to partial alignments with *R*^*2*^ values of 0.9 and 0.5, respectively. Together, the anchor alignment algorithm can avoid erroneous counting and can also distinguish mature miRNA reads from precursor miRNA reads accurately.

### 3.2 Anchor optimization

To include sncRNA variants that bear small overhangs or internal insertions/deletions/mutations in the sncRNA-Seq reads, we tested a number of anchor sequences to see which ones gave the best alignment results. We first tested two 5nt anchors by aligning the simulation datasets against the reference sncRNA datasets downloaded from various sncRNA databases (Figure 2A). The simulation dataset containing all the known sncRNAs aligned perfectly to the sncRNA reference datasets (*R*^*2*^=1). However, when the simulation datasets containing 1-2nt overhangs at either end were used, only partial alignment (*R*^*2*^=0.87) was achieved due to the gap-opening penalty caused by those miRNA variants (Figure 2A, Supplementary file 1: Figure S1). Since these miRNA variants are likely synthesized by the cell and the 1-2nt mutations are probably due to sequencing errors, they should not be excluded from annotation. To accommodate theses sncRNA variants, we designed C/G repeat anchors of different lengths (5nt for the reads and 10nt for the references) based on the fact that C and G are the least common nucleotides at the ends of miRNAs and thus, can have higher specificity (Supplementary file 1: Figure S2). Using C/G repeat anchors for alignment, a 1-2nt overhang in the read sequences would lead to a mismatch instead of a gap-opening penalty, which allows for inclusion of these sncRNA variants into the counts, leading to an increased alignment rate (*R*^*2*^ from 0.87 to 0.92) (Figure 2A, Supplementary file 1: Figure S1). We also examined the AG anchors as well as other possible single nucleotide anchors, and found that anchors with the C/G combination consistently yielded the highest alignment rates (Supplementary file 1: Figure S2). We also evaluated different anchor lengths (5-10nt), and the 5nt C/G anchors were chosen as the default setting for alignment analyses using AASRA, based on the better performance compared to other lengths (Supplementary file 1: Figure S3). By fine-tuning the parameters of AASRA, the optimal setting was determined such that the sncRNA variants with 1-2nt overhangs, internal insertions/deletions/mutations, could be included into the final counts (Supplementary file 1: Figure S4). For annotating sncRNA sequencing reads containing small internal insertions/deletions/mutations, AASRA (with the use of the C/G anchors) consistently outperformed the Bowtie2-based direct sncRNA-sncRNA mapping method (Supplementary file 1: Figure S5). Overall, these data indicate that the C/G anchor-based alignment algorithm of AASRA allows for efficient mapping of not only perfect-matching sequencing reads, but also reads with small (1-2nt) overhangs and internal insertions, deletions or mutations.

**Figure 2.**
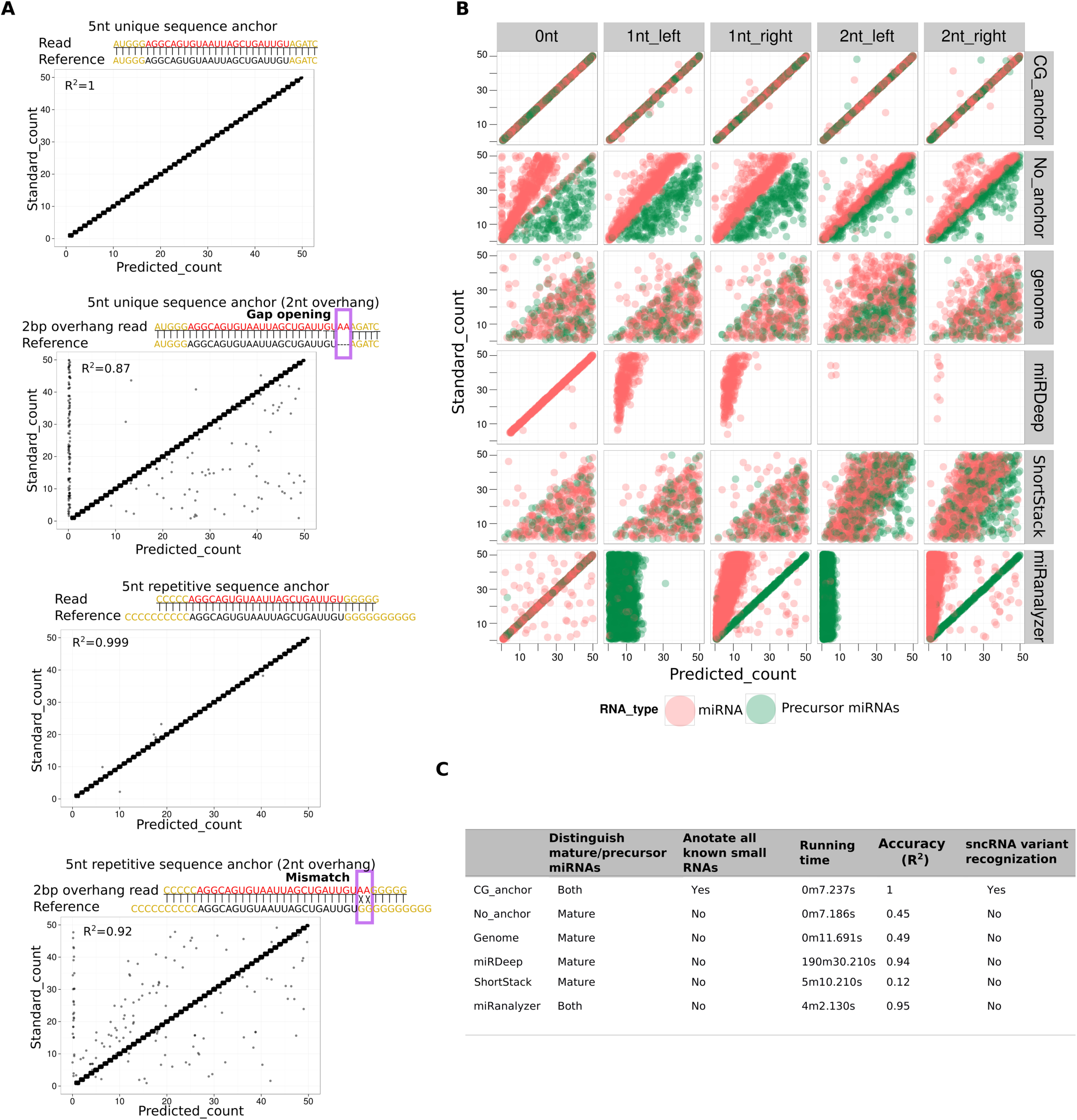
Anchor optimization and performance comparison between AASRA and three existing sncRNA annotation pipelines. (A) C/G anchors outperformed other anchors because the C/G anchors could turn the gap-opening penalty (causing exclusion) into mismatch penalty (leading to inclusion). The use of a non-C/G anchors could align the simulation data without overhangs perfectly (*R*^*2*^=1), but simulation sequences with 2nt overhangs were only aligned partially (*R*^*2*^=0.87) due to gap-opening penalty that excluded many miRNA variants. In contrast, the use of C/G anchors aligned simulation datasets with or without 2nt overhangs almost perfectly (*R*^*2*^= 0.999 and 0.92, respectively) because those 2nt overhangs were treated as mismatches rather than gaps and thus, those variants were counted and annotated. (B) Performance comparison between AASRA and three existing sncRNA annotation software packages (miRDeep, ShortStack and miRanalyzer). Simulation datasets containing both mature (red dots) and premature (green dots) miRNA sequences with 0-2nt overhangs were aligned to the reference sncRNA dataset using direct sncRNA-sncRNA alignment (no-anchor), direct alignment to the genome (genome), miRDeep, ShortStack and miRanalyzer. (C) Summary of the performance of AASRA and other five sncRNA annotation pipelines tested.

### 3.3 Performance comparison between AASRA and three existing sncRNA annotation software packages

To demonstrate the superior performance of AASRA, we generated simulation datasets containing mature and precursor miRNAs with 0, 1-2nt overhangs at either end, and annotated the simulation sequence reads against the reference miRNA datasets downloaded from the miRBase using AASRA and three popular software packages for miRNA annotation, including ShortStack [17], miRDeep [19] and miRanalyzer [18] (Figure 2B). The simulation sequences were aligned almost perfectly to the references datasets using AASRA for both mature and precursor miRNAs with or without overhangs (*R*^*2*^ ≈ 1) (Figure 2B, 2C). In contrast, direct Bowtie2-based mapping of the simulation miRNA and precursor miRNA sequences with or without overhangs to the reference miRNA datasets or to the mouse genome resulted in poor alignment rates (*R*^*2*^ = 0.45 −0.49). Although miRDeep could map sequences perfectly matching the known mature miRNAs efficiently (*R*^*2*^ = 0.94), it failed to align either precursor miRNA sequences or mature miRNA sequences with overhangs (Figure 2B, 2C), largely due to its strict length control criteria [19]. Thus, miRDeep cannot annotate precursor miRNAs, mature miRNAs with mismatches, or other sncRNAs with staggered sequence patterns (e.g., piRNAs, mitosRNAs, tsRNAs, etc.). ShortStack, similar to the direct genome alignment method, could only annotate a small fraction of the simulation sequences, largely due to repetitive and ambiguous counting. miRanalyzer utilizes a three-phase alignment procedure (i.e., mature miRNA alignment → pre-miRNA alignment → genome alignments) in conjunction with length control. miRanalyzer annotated the simulation data without overhangs as efficiently as AASRA (*R*^*2*^= 0.95), but failed to annotate simulation data containing overhangs because it dose not tolerate mismatches. In summary, AASRA appeared to be ideal for annotating known sncRNA species simultaneously with the capability of distinguishing mature and precursor miRNAs, and recognizing sncRNA variants with small overhangs and/or internal insertions/deletions, with a speed faster than any of the five pipelines tested (Figure 2C).

### 3.4 AASRA-based annotation of sperm sncRNAs

Two advantages of AASRA over the existing sncRNA annotation software packages include the following: 1) it can identify novel sncRNA variants with small overhangs or internal insertions, deletions or mutations. 2) It can annotate not only miRNAs (both mature miRNAs and pre-miRNAs), but also all known sncRNA species collected in various databases. A key question remains: do those sncRNA variants exist in the sncRNA–Seq reads by a substantial proportion? If so, these sncRNA variants should not be overlooked in quantitative analyses. To answer this question, we annotated the sperm sncRNA-Seq data generated by both the Ion Proton and the Illumina sequencers using both AASRA and miRDeep. AASRA simultaneously annotated nine known species of sncRNAs from mouse sperm sncRNA-Seq reads (Figure 3A). By comparing the unique mature miRNA counts determined by miRDeep and AASRA, we found that AASRA identified 37% more unique mature miRNA counts than miRDeep (Figure 3B). While miRDeep could not annotate precursor miRNAs, AASRA identified both mature and precursor miRNAs (Figure 3C). Interestingly, murine sperm appeared to contain numerous precursor miRNAs, which would not have been identified using miRDeep or other sncRNA annotation software packages (Figure 3C). Further examination of the alignment results for the four miRNAs (mir-376a, mir-361, mir-93 and mir-4660) revealed that AASRA not only identified more mature miRNAs than miRDeep, but also detected various miRNA variants, including those containing small (1-2nt) overhangs, internal insertions, deletions or mutations, whereas these sncRNA variants were not detected by miRDeep (Figure 3D). For example, ∼80% of the sequencing reads aligned to miR-93 all contained overhangs, which could be either biological variants of miR-93 or sequencing errors. Regardless, such a large number of miR-93 variants would have been totally ignored if other existing software packages were used (Figure 3D). If one wants to exclude these sncRNA variants, a more stringent alignment can be performed through adjusting the parameters, including anchor sequence and mismatch penalty. For example, four levels of specificity settings (high_specificity1, 2, 3 and ultra) (Supplementary file 1: Figure S6A) were tested for sequence alignment stringency. At the ultra-high specificity setting, AASRA could eliminate all the sequences with 1-2nt overhangs in the simulation data (Supplementary file 1: Figure S6B). Under the same setting, perfectly-matched miRNAs could be readily identified from a mixture of miRNA sequences with 1-2nt overhangs (Supplementary file 1: Figure S6C). The ultra-high specificity setting made AASRA function similarly as miRDeep, whereas a less stringent setting allowed for identification of miRNA variants (Supplementary file 1: Figure S6D). It will be up to the investigators to decide whether those sncRNA variants should be included or excluded in the final counts during sncRNA annotation depending on the nature of specific experiments conducted.

**Figure 3.**
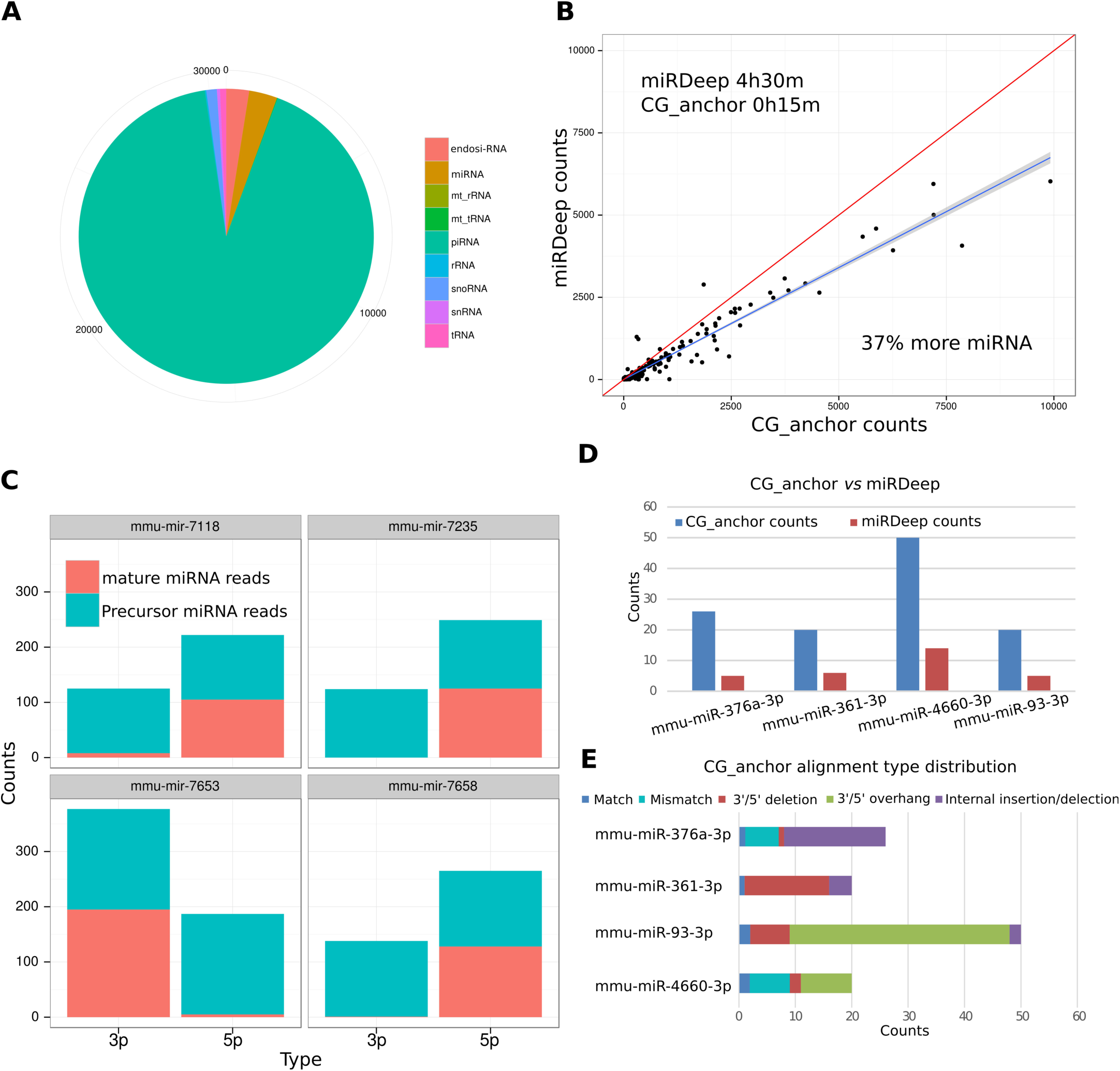
Annotation of sperm sncRNA-Seq data using AASRA. (A) Pie chart showing the count distribution of nine sncRNA species in murine sperm annotated using AASRA. (B) Scatter plot showing that AASRA could identify 37% more miRNAs than miRDeep in 1/12 of the time needed by miRDeep. (C) Counts of four miRNAs and their precursors in the sperm sncRNA-Seq data, as determined by AASRA. (D) Counts of four mature miRNAs in murine sperm sncRNA data, as determined by AASAR and miRDeep.(E) The contents of the AASRA counts of the four mature miRNAs shown in Figure D. Note that mismatches, deletions, insertions, and overhangs appear to be common in the sncRNA sequencing reads.

## 4. Discussion

The rapid advance of next-gen sequencing technologies has led to the discovery of hundreds thousands of sncRNAs [2]. Increasing lines of evidence suggest that these sncRNAs play regulatory roles critical to development and physiology [2]. Despite the rapid pace of sncRNA discovery, the bioinformatic tools for sncRNA annotation are very limited. None of the currently available sncRNA annotation pipelines can annotate simultaneously all known sncRNA species, nor can they tolerate sequences with mismatches although these sncRNA variants are likely due to sequencing errors, but biologically relevant. AASRA utilizes a unique, anchor alignment-based algorithm, and is capable of annotating all known sncRNAs simultaneously. The specificity setting of AASR is adjustable such that small mismatches due to overhangs, insertions, deletions, or mutation, can be either included or excluded. AASRA can identify a much greater number of sncRNA counts (e.g., ∼37% more identified from the murine sperm sncRNA-Seq data) compared to any of the existing pipelines because of the use of the anchor alignment algorithm. This feature offers the possibility of minimizing quantification bias caused by 1) over-counting (due to double and ambiguous alignments), and/or 2) exclusion of variant sequences in the sncRNA-Seq data (although these variants should be counted because they are produced by the cells, but simply slightly different from the main sncRNA sequences most likely due to sequencing errors). The fact that these variant sequences account for a large proportion of the total counts (e.g., up to 80% for mmu-miR-93), elimination of these variants would greatly skew the real expression profile, leading to inaccurate interpretation and conclusions. Since all existing sncRNA annotation software packages do not have these functions, AASRA will be very useful for investigators to revisit their sncRNA data to see how many variants were inadvertently excluded, and whether such exclusion had caused quantitation bias that would compromise their conclusions. Depending on the needs of the investigators, those variants can also be excluded by applying more strict alignment parameters.

The capability to annotate the precursor miRNAs is another useful feature of AASRA. Interestingly, a large number of precursor miRNAs appear to be present in sperm, which would not have been discovered if other existing programs were used. Although miRanalyzer can annotate precursor miRNAs, it can only annotate those with perfect matches, and those with small overhangs or minor mismatches would be ignored. Mature miRNAs have been found in sperm of multiple species, including mouse [40, 42], rat [33, 43], cow [44], horse [45], monkey [40, 46] and human [46, 47]. However, sperm-borne precursor miRNAs have not been reported. Given that these precursor miRNAs can be potentially delivered into the eggs during fertilization, their potential regulatory roles would be an intriguing topic for future investigation.

In summary, AASRA represents the first universal sncRNA annotation software package, which allows for simultaneous annotation of all known sncRNAs with high speed and accuracy. AASRA can annotate not only known sncRNA species, but also sncRNA variants containing small overhangs, or internal deletions/insertions/mutations. AASRA provides another useful bioinformatic tool for studying sncRNA biology.

## Authors’ contributions

CT and WY conceived and designed the study, CT wrote the software with the assistance from YX, CT and WY wrote the manuscript. All authors read and approved the final manuscript.

## Acknowledgements

The authors would like to thank the rest of the Yan lab for helpful discussion during the development of AASRA.

## Supplementary file 1

**Figure S1.** Comparison of alignment accuracy between AASRA (CG_anchor) and the direct sncRNA-sncRNA alignment method (No_anchor) using sncRNA simulation data containing miRNAs, endo-siRNAs, piRNAs, snRNAs and tRNAs. **Figure S2.** Comparison of alignment accuracy between the CG or AG anchor using miRNA simulation data containing mature and premature miRNAs. **Figure S3**. Comparison of alignment accuracy among anchors with different lengths (5-10nt) using sncRNA simulation datasets containing miRNAs, endo-siRNAs, piRNAs, snRNAs and tRNAs with (0nt) or without 1-2nt overhangs. **Figure S4.** Comparison of alignment accuracy affected by different Bowtie2 parameters using sncRNA simulation datasets containing miRNAs, endo-siRNAs, piRNAs, snRNAs and tRNAs with (0nt) or without 1-2nt overhangs. **Figure S5.** Comparison of alignment accuracy between AASRA (CG_anchor) and the direct sncRNA alignment method (No_anchor) using sncRNA simulation datasets containing miRNAs, endo-siRNAs, piRNAs, snRNAs and tRNAs with 1nt internal deletions (A), insertions (B) or mutations (C). **Figure S6.** The AASRA ultra-high specificity function.

## Supplemental Information

**Figure S1.**
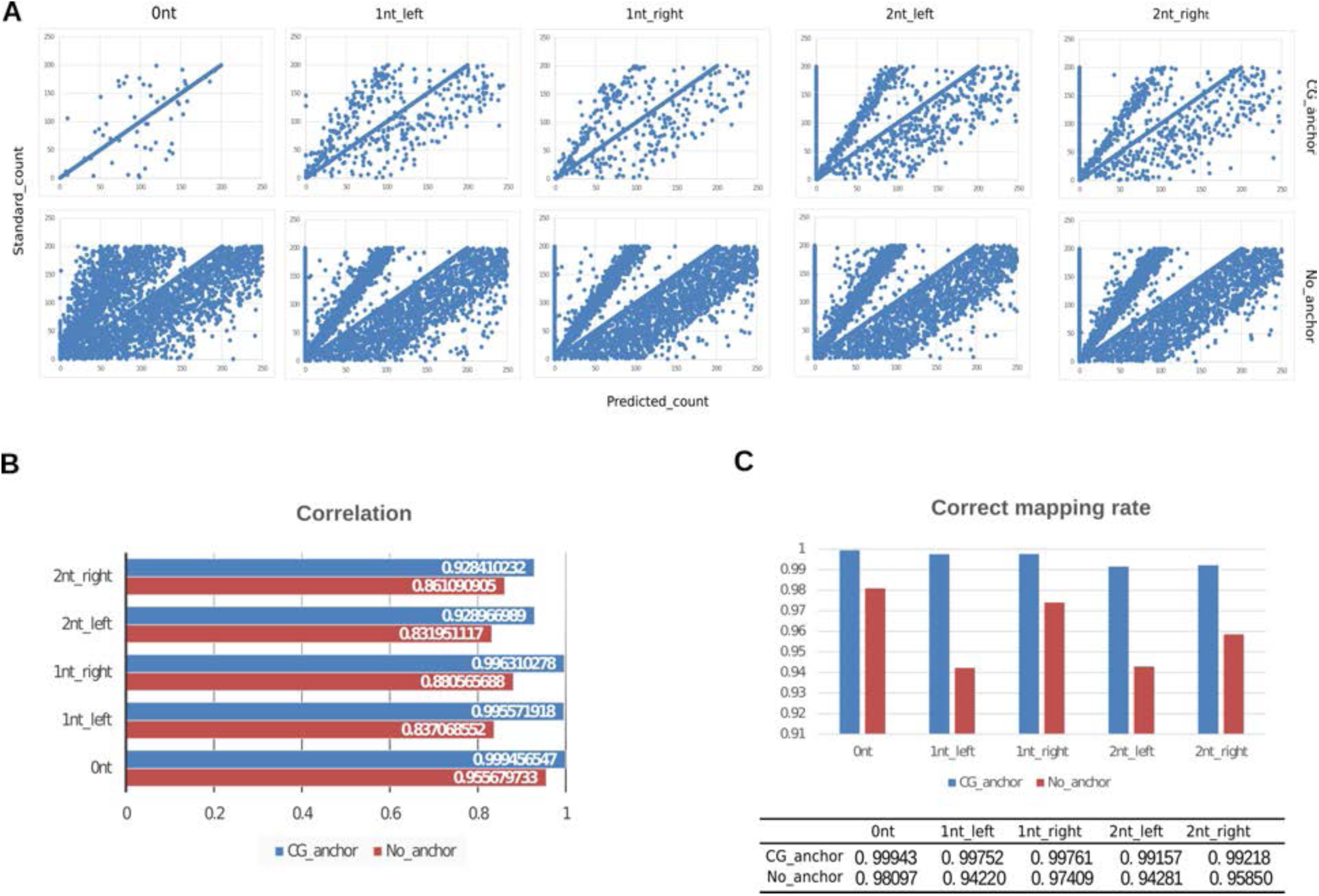
Comparison of alignment accuracy between AASRA (CG_anchor) and the direct sncRNA-sncRNA alignment method (No_anchor) using sncRNA simulation data containing miRNAs, endo-siRNAs, piRNAs, snRNAs and tRNAs. (**A**) Scatter plots showing alignment of the simulation sncRNA datasets with or without 1-2nt overhangs by AASRA (CG_anchor) and the direct sncRNA alignment method. (**B**) Bar graphs comparing the correlation coefficient (*R*^*2*^) values between predicted counts (calculated by the algorithm) and standard counts (known for the simulation data) identified using AASRA (CG_anchor) and the direct sncRNA alignment method (No_anchor). (**C**) Bar graphs comparing the correct mapping rates of the simulation datasets between AASRA and the direct sncRNA alignment. The correct mapping rate is defined as the number of correctly mapped reads/ total reads.

**Figure S2.**
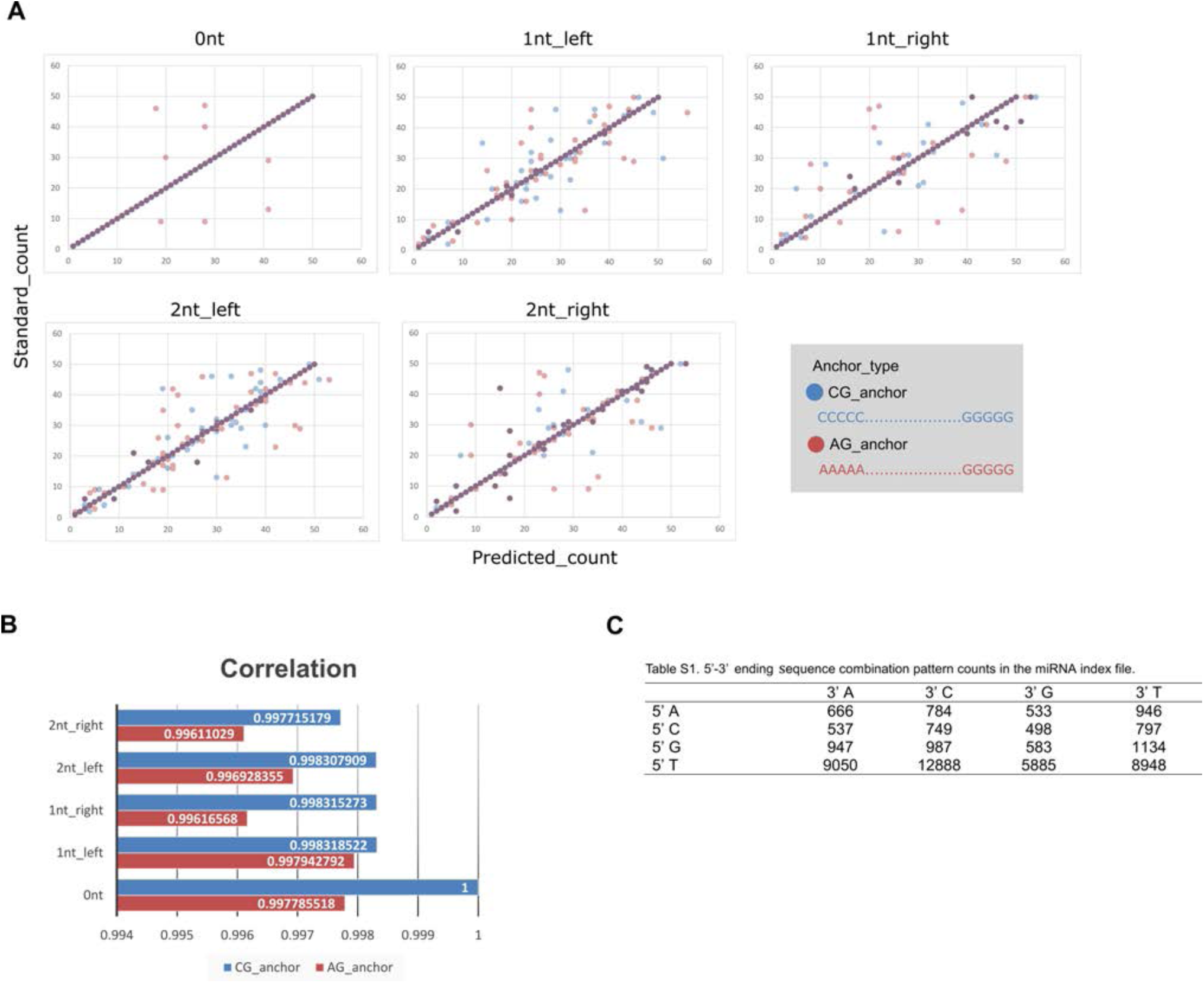
Comparison of alignment accuracy between the CG or AG anchor using miRNA simulation data containing mature and premature miRNAs. (**A**) Scatter plots showing correlations between the predicted counts (calculated by the algorithm) and standard counts (known for the simulation data) derived from alignment using the CG anchor (blue dots) or the AG anchor (red dots). The CG anchor yielded better results than the AG anchor. (**B**) Bar graphs comparing the correlation coefficient values between the predicted counts (calculated by the algorithm) and standard counts (known for the simulation data) identified using the CG or AG anchor for alignment. (**C**) Frequency of the four nucleotides at both ends of miRNAs in the miRNA index file. MiRNA sequences that start with cytosine and end with guanine are the least common and thus, the CG anchor is a better choice due to a lower frequency at both ends of sncRNAs.

**Figure S3.**
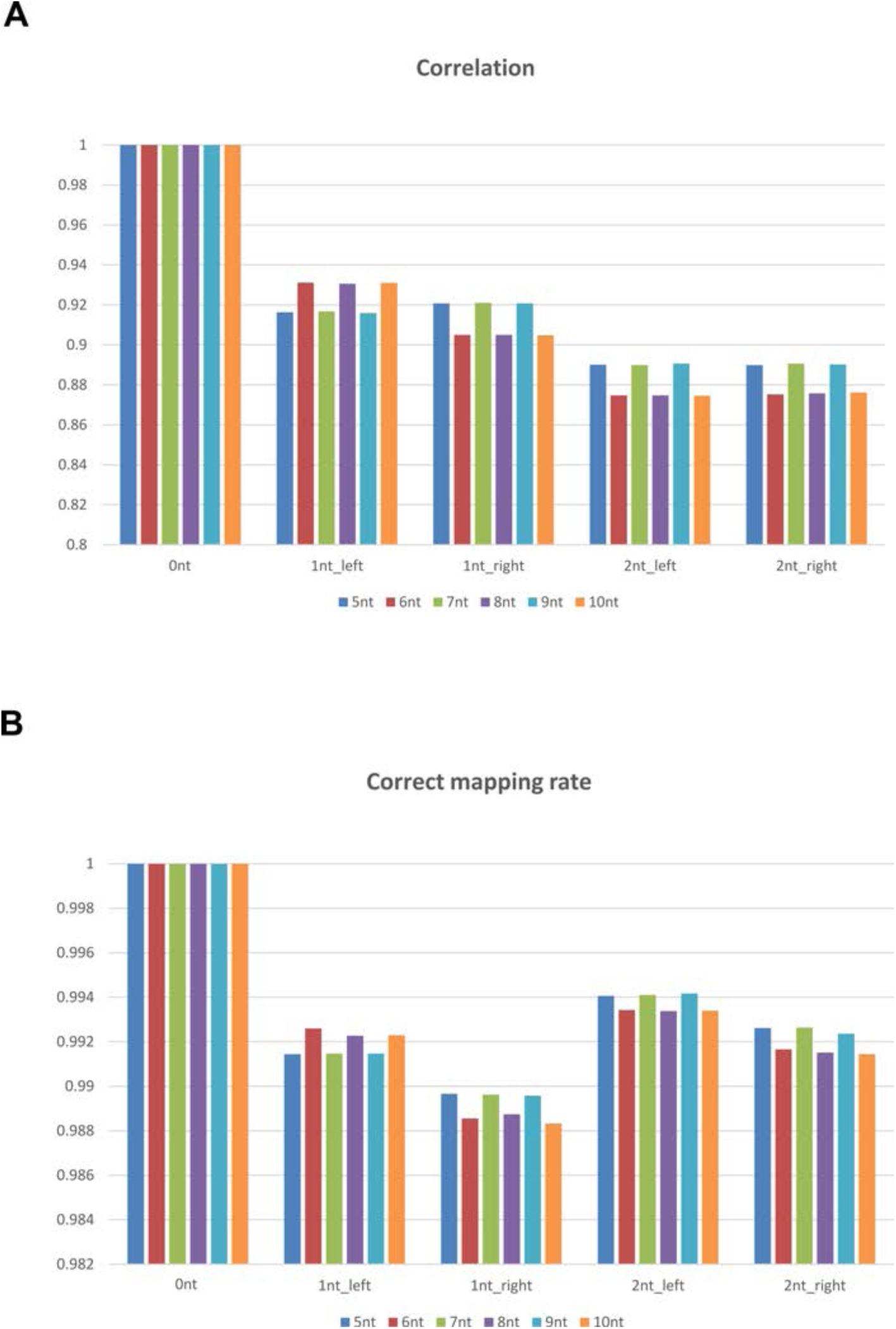
Comparison of alignment accuracy among anchors with different lengths (5-10nt) using sncRNA simulation datasets containing miRNAs, endo-siRNAs, piRNAs, snRNAs and tRNAs with (0nt) or without 1-2nt overhangs. (**A**) Bar graphs showing correlation rates between predicted counts (by software calculation) and standard counts (simulation data) when anchors of different lengths (5-10nt) were used for alignment. (**B**) Bar graphs showing correct mapping rates of the simulation datasets when anchors of different lengths (5-10nt) were used for alignment.

**Figure S4.**
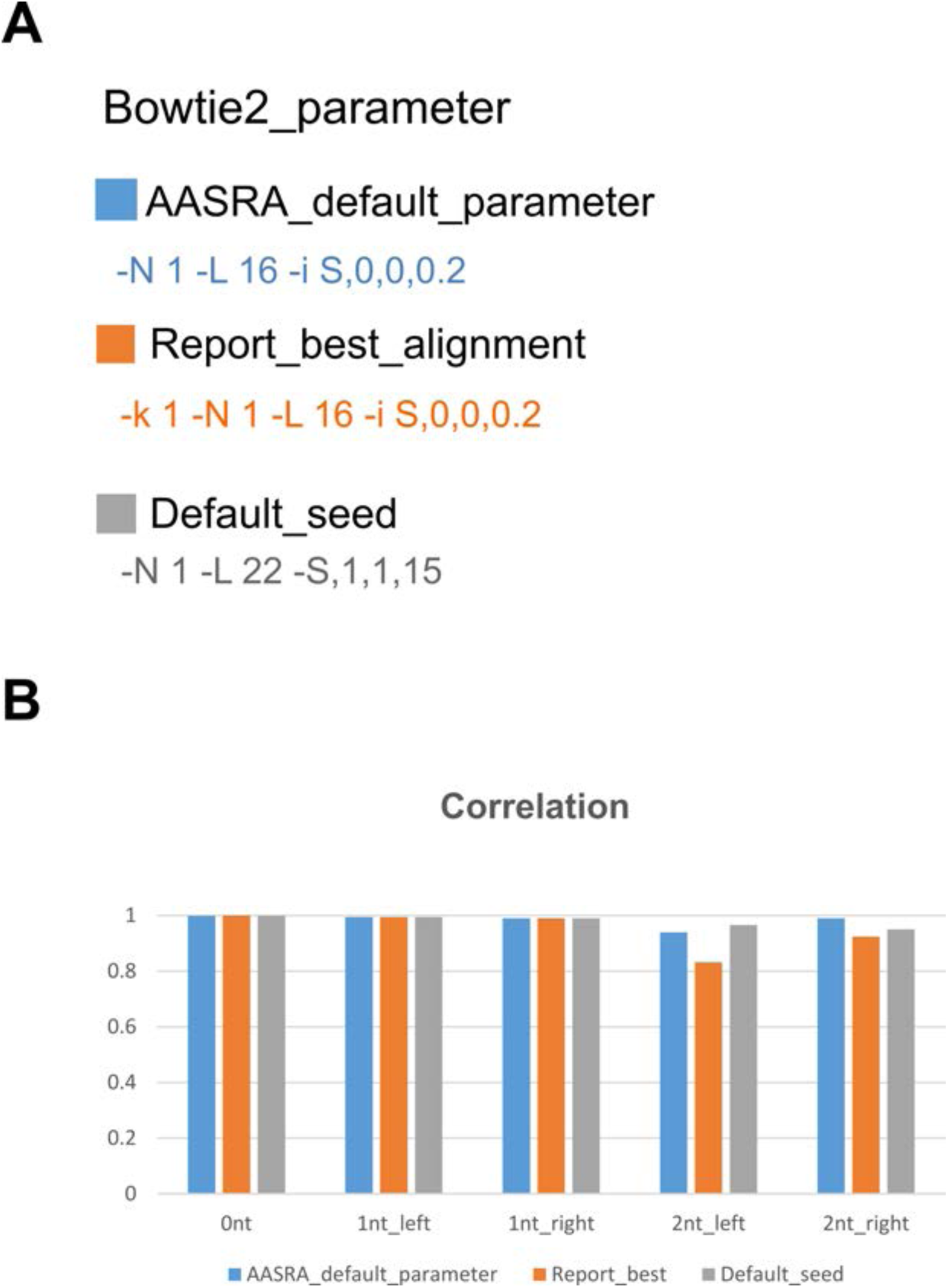
Comparison of alignment accuracy affected by different Bowtie2 parameters using sncRNA simulation datasets containing miRNAs, endo-siRNAs, piRNAs, snRNAs and tRNAs with (0nt) or without 1-2nt overhangs. (**A**) Bowtie2 commands with parameters used for comparison. AASRA default parameters were optimized based on those yielding the best-reported alignment results by Bowtie2 (Reprot_best_aligment). Bowtie default parameters (Default_seed) are tested as well. Seed length (-L) was optimized for miRNA alignment in AASRA. AASRA default allows 1 mismatch in seed sequence alignment (-N 1). (**B**) Bar graphs showing the correlation rates between predicted counts (by software calculation) and standard counts (simulation data) when 3 different Bowtie2 parameters, as illustrated in panel A, were used for alignment. Default AASRA parameter produces the best overall alignment accuracy for simulation datasets.

**Figure S5.**
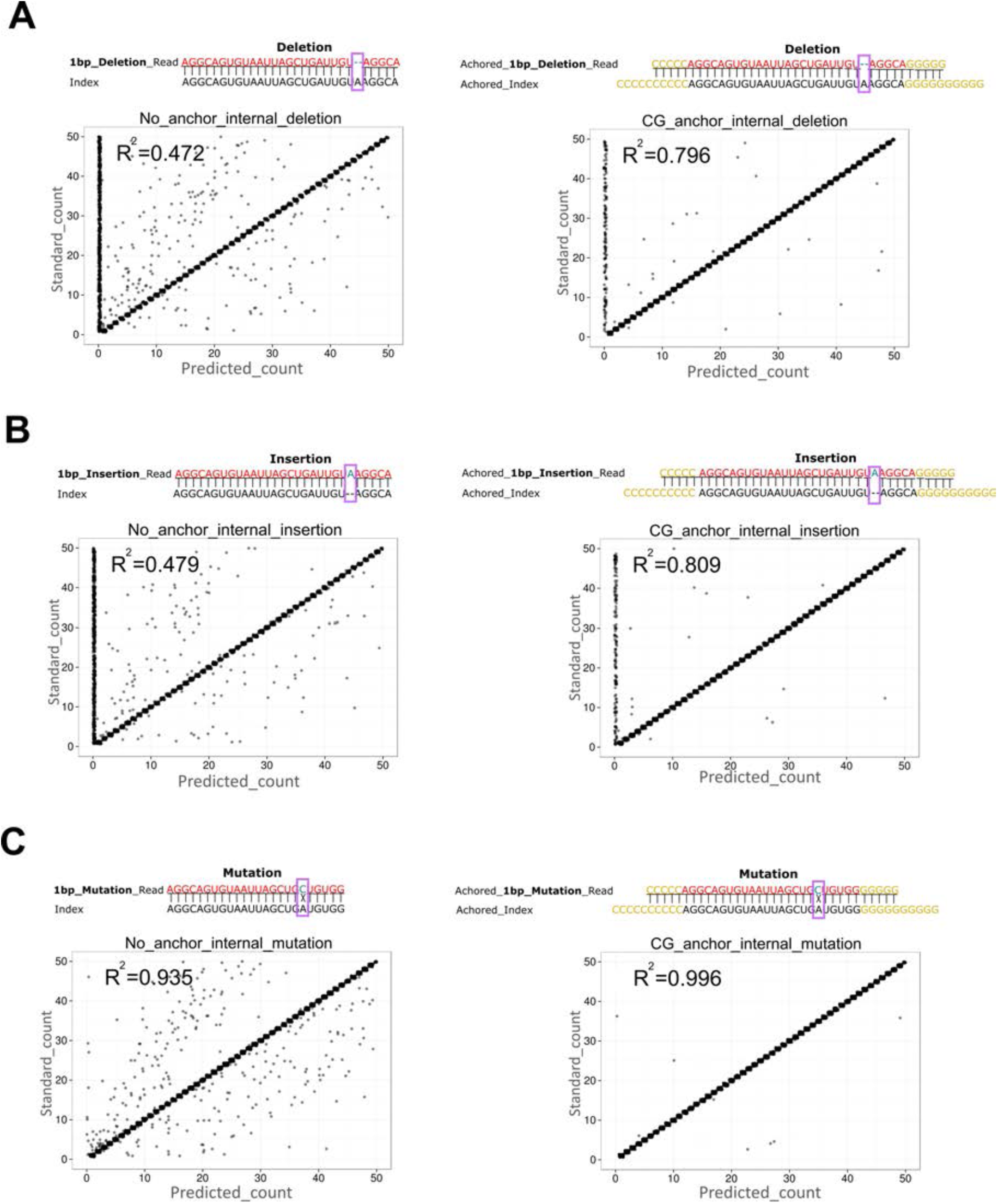
Comparison of alignment accuracy between AASRA (CG_anchor) and the direct sncRNA alignment method (No_anchor) using sncRNA simulation datasets containing miRNAs, endo-siRNAs, piRNAs, snRNAs and tRNAs with 1nt internal deletions (A), insertions (B) or mutations (C).

**Figure S6.**
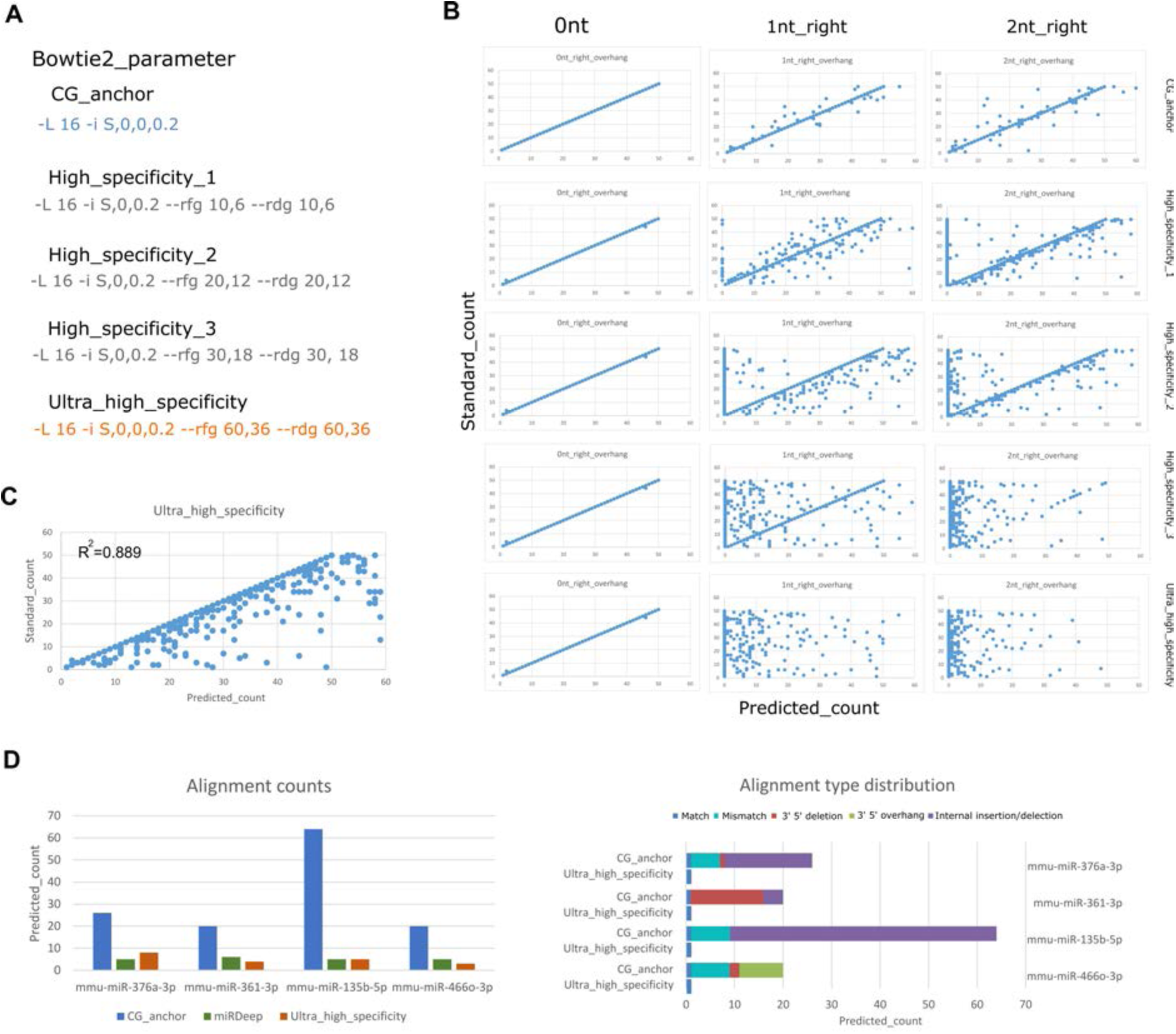
The AASRA ultra-high specificity function. (**A**) The parameters adjusted for various levels of alignment stringency. -L is the seed length of alignment; –i functions to govern the interval between two seed substrings used during multispeed alignment, controlling sensitivity and speed; --rfg --rdg controls the gap opening penalty. The highest stringency (ultra_high_specificity) setting can eliminate sequences containing 1-2nt overhangs. (**B**) Scatter plots showing alignment results at the four levels of specificity settings (High_specificty_1-3, and ultra_high_specificity) using sncRNA simulation datasets containing miRNAs, endo-siRNAs, piRNAs, snRNAs and tRNAs with (0nt) or without 1-2nt overhangs. (**C**) No effects of sequences with overhangs on the alignment results under the ultra-high specificity setting. Simulation datasets containing no (0nt) or 1-2nt overhangs were merged and used for mapping against the reference dataset. The ultra_high_specificity setting effectively eliminated the interference from the reads with overhang nucleotides (*R*^*2*^=0.889). (**D**) Bar graphs showing the counts of sperm sncRNA reads aligned to four mature miRNAs using miRDeep and AASRA at default (CG_anchor) and ultra-high specificity settings (Ultra_high_specificity). Counts for different variant types of the four miRNAs identified using AASRA under the default (GC_anchor) and ultra high specificity (Ultra_high_specificity) settings. The ultra_high_specificity setting effectively removed miRNA variants in the sncRNA-Seq data.

